# Fitness Benefits of Loss of Heterozygosity in Saccharomyces *Hybrids*

**DOI:** 10.1101/452748

**Authors:** Samuel M. Lancaster, Celia Payen, Caiti Smukowski Heil, Maitreya J. Dunham

**Affiliations:** Department of Genome Sciences, University of Washington, Seattle, WA; Current Address: Department of Genetics, Stanford University, Stanford, CA; Current Address: Industrial Biosciences DuPont, Wilmington, DE

**Author notes:** Corresponding Author: Maitreya Dunham, Foege Building S403B, Box 355065; 3720 15th Ave NE; Seattle WA 98195-5065.

**Keywords:** hybrid, yeast, loss of heterozygosity, adaptation

## Abstract

With two genomes in the same organism, interspecific hybrids have unique opportunities and costs. In both plants and yeasts, wild, pathogenic, and domesticated hybrids may eliminate portions of one parental genome, a phenomenon known as loss of heterozygosity (LOH). Laboratory evolution of hybrid yeast recapitulates these results, with LOH occurring in just a few hundred generations of propagation. In this study, we systematically looked for alleles that are beneficial when lost in order to determine how prevalent this mode of adaptation may be, and to determine candidate loci that might underlie the benefits of larger-scale chromosome rearrangements. These aims were accomplished by mating *Saccharomyces uvarum* with the *S. cerevisiae* deletion collection to create hybrids, such that each nonessential *S. cerevisiae* allele is deleted. Competitive fitness assays of these pooled, barcoded, hemizygous strains, and accompanying controls, revealed a large number of loci for which LOH is beneficial. We found that the fitness effects of hemizygosity are dependent on the species context, the selective environment, and the species origin of the deleted allele. Further, we found that hybrids have a larger distribution of fitness consequences vs. matched *S. cerevisiae* hemizygous diploids. Our results suggest that LOH can be a successful strategy for adaptation of hybrids to new environments, and we identify candidate loci that drive the chromosomal rearrangements observed in evolution of yeast hybrids.

## INTRODUCTION

Hybrid organisms are common in nature, particularly in fungi and plants where an estimated 4% of flowering plants and 7% of ferns are hybrids (Otto and Whitton 2000). Even the human genome is now recognized to contain substantial introgressions – remnants of ancient hybridization – that are thought to be adaptive (Huerta-Sanchez et al. 2014; Dannemann et al. 2016; Gittelman et al. 2016; Racimo et al. 2017). Hybrids have been created via artificial selection in agriculture, industry, and the laboratory. For example, wheat, a pillar of civilization, is a triple hybrid between three grass species (Brenchley et al. 2012). Hybridization and introgression are also abundant in budding yeast (reviewed in Morales and Dujon), where hybrids have been found to possess adaptive advantages over their parental species (e.g., Stelkens et al. 2014), show desirable properties as industrial organisms (e.g., Mertens et al. 2015; Peris et al. 2017) and contribute to the emergence of fungal pathogens (Morales and Dujon 2012; Pryszcz et al. 2015; Schroder et al. 2016; Mixao and Gabaldon 2018). A whole genome duplication ancestral to *Saccharomyces* yeasts – a defining characteristic of the clade – has been recognized as a hybridization event (Marcet-Houben and Gabaldon 2015). Since *Saccharomyces* has relatively weak prezygotic barriers to speciation (Maclean and Greig 2008; Murphy and Zeyl 2012), *Saccharomyces* is particularly rife with hybridization, and includes hybridization between species as distant as 20 million years diverged (~80% amino acid and nucleotide identity), which are capable of intermating (Martini and Martini 1987; Naumova et al. 2005; Dunn and Sherlock 2008; Muller and McCusker 2009; Libkind et al. 2011; Nguyen et al. 2011; Almeida et al. 2014; Perez-Traves et al. 2014). Two common yeasts that originated as hybrids between *S. cerevisiae* and cryotolerant species have even received designation as hybrid species: the wine yeast *S. bayanus,* a triple hybrid between *S. cerevisiae, S. uvarum*, and *S. eubayanus* (Gonzalez et al. 2006; Sipiczki 2008); and the lager yeast *S. pastorianus,* which is a hybrid between *S. cerevisiae* and *S. eubayanus* (Martini and Martini 1987; Dunn and Sherlock 2008; Libkind et al. 2011; Nguyen et al. 2011; Perez-Traves et al. 2014). These species highlight the observation that fermentation environments are particularly rich in hybrids, spanning genera including *Saccharomyces, Dekkara,* and *Pichia* (Borneman et al. 2014; Smukowski Heil et al. 2018a).

Similar to plant hybrids (reviewed in Chester et al. 2010), yeast hybrids can shed large portions of their genomes from one or both species during evolution (Otto and Whitton 2000; Sun and Xu 2009; Chester et al. 2010; Csoma et al. 2010; Louis et al. 2012; Peris et al. 2012; Pryszcz et al. 2015; Chen et al. 2017; Emery et al. 2018). Resolution of the ancestral whole genome duplication in *Saccharomyces* involved loss of the majority of duplicated genes, in a process that began shortly after the initial hybridization event (Scannell et al. 2007). These large-scale changes in genome structure and content have been recapitulated in part in the laboratory, demonstrating the rapidity with which these changes can occur and confirming their potential to contribution to adaptation. For example, experimental evolution of yeast hybrids under a number of selective conditions found genome rearrangements after only a few hundred generations (Kunicka-Styczynska and Rajkowska 2011; Piotrowski et al. 2012; Sanchez et al. 2017a; Smukowski Heil et al. 2017). The genome regions affected are dependent on the selective pressure used, and observed events include whole chromosome aneuploidy, both focal and chromosome arm amplifications, translocations, gene fusions, and LOH. Our previous work demonstrated that LOH can result from selection on one species allele and loss of the other. Using a candidate gene approach, we identified a single gene (*PHO84*) whose allelic differences explained the majority of the fitness benefit in evolved populations relative to their ancestor. However, additional, as yet unidentified driver genes must exist to fully account for the evolved strains’ fitness improvements, and many observed LOH regions remain unexplained. More broadly, the degree to which LOH is a product of genetic drift versus selection is not yet clear. To further complicate matters, improved fitness caused by LOH could have a number of possible explanations, including selecting for the better species’ alleles, uncovering beneficial recessive alleles, and/or resolving hybrid incompatibilities. Studying LOH in hybrid yeast allows a systematic approach that facilitates insights into these phenomena and provides a foundation to guide further investigation into hybrid biology in other, less tractable, contexts.

Such systematic approaches have been made possible via the creation of genome-scale deletion collections, including a near-comprehensive set of diploid *S. cerevisiae* strains hemizygous for every gene (Giaever et al. 2002; Deutschbauer and Davis 2005). These strains were created such that each carries a unique DNA barcode, facilitating pooled assays for competitive growth in a variety of conditions. Many studies have illustrated that heterozygous deletions can cause fitness defects (“haploinsufficiency”), and a smaller number have also found fitness increases, or haploproficiency (Delneri et al. 2008; Pir et al. 2012; Ohnuki and Ohya 2018). Previously, in order to determine driver mutations, our lab identified haploinsufficient and haploproficient loci in the deletion collection in environments matching laboratory evolution studies (Payen et al. 2016). However, since these loci were identified in *S. cerevisiae* diploids, the degree to which they explain the prevalence of, and genetic drivers for, hemizygosity in hybrids is unclear. There is reason to believe that loci important for hybrid adaptation are likely to differ from those important in purebred diploids. For example, Herbst et al., 2017, found that in *S. paradoxus* x *S. cerevisiae* hybrids hundreds of allelic deletions decreased the growth rate of hybrids but not of *S. cerevisiae* diploids.

In order to understand hybrid LOH, in this study we utilized two divergent species: *S. cerevisiae* and *S. uvarum.* We previously evolved hybrids and diploids of these species in nutrient-limited chemostat culture (Gresham et al. 2008; Sanchez et al. 2017a; Smukowski Heil et al. 2017; 2018b). We created thousands of *S. cerevisiae* x *S. uvarum* hybrid yeast strains by mating the *S. cerevisiae* nonessential deletion collection to WT *S. uvarum*. These collections of hybrid yeast, along with control populations, were assayed for competitive fitness in three nutrient-limited environments matched to our previous evolution conditions. We find that haploproficiency is common, and more loci are haploproficient in hybrids than in *S. cerevisiae* diploids, providing giving them more opportunities to adapt via this mechanism. Specific haploproficient loci rarely overlap between *S. cerevisiae* diploids and interspecific hybrids, indicating that simple gene dosage changes are unlikely to explain their adaptive benefit, and/or that dosage sensitivity is strongly dependent on species context. Furthermore, the specific loci are largely private to single selection environments, agreeing with our previous experimental evolution findings showing repeated occurrence of LOH for specific regions was also condition-specific. Finally, fitness effects are allele-specific – fitness consequences of deletion of the *S. cerevisiae* allele in the hybrid had no correlation with the fitness consequences of deletion of the *S. uvarum* allele. Again, these results are consistent with prior observations of species preference for LOH in evolved hybrids. They argue that if relief of genetic incompatibilities is a relevant mechanistic explanation for adaptation, then such incompatibilities must be acting in an allele-specific manner. Our study demonstrates that hybrids offer a unique fitness landscape with potentially more beneficial mutations, which may contribute to their unique ability to adapt, and it provides attractive candidate genes for future study.

## RESULTS

We sought to discover the genome-wide fitness effects of hemizygosity in hybrid *Saccharomyces* in three nutrient-limited conditions that correspond with those previously used for experimental evolution. To this end, we mated *S. uvarum* to the *S. cerevisiae* haploid deletion collection creating thousands of hybrid yeast strains, each with a *S. cerevisiae* allele deleted. For comparison, we used a matched collection of *S. cerevisiae* hemizygous deletion strains. Additionally, we created control collections of thousands of WT *S. cerevisiae* and WT hybrid strains that contain unique DNA barcodes but are otherwise isogenic. (All collections are described in Table 1.) These control libraries allowed us to empirically measure technical and biological variation in our strain construction, growth, and sequencing pipeline. All strains were assayed for relative fitness via pooled competitive growth for 25 generations in glucose, phosphate, and sulfate limited chemostat culture followed by barcode sequencing. The barcodes counts track strain abundance over time, allowing us to derive competitive fitness scores (see Materials and Methods, Supplemental Table S1). Each experiment was performed in biological replicate (Supplemental Fig. S1). We confirmed that this pooled approach accurately reflects strain fitness by comparing the results to pairwise competitions of individual deletion strains vs. a GFP-marked WT competitor (Supplemental Fig. S2, Supplemental Table S2).

**Table 1.** Strain collections and datasets used in this study.

| Collection | Description |
| --- | --- |
| *S. cerevisiae* Hemizygous Deletion Collection (“purebred” collection) | *S. cerevisiae* hemizygous deletion collection (from Giaever et al., 2002). Fitness data from (Payen et al 2016). |
| Hybrid Hemizygous Deletion Collection | *S. cerevisiae* haploid deletion collection (from Giaever et al., 2002) mated to WT *S. uvarum*. |
| *S. cerevisiae* Barcoded Control Library | Collection of barcoded WT haploid *S. cerevisiae* strains (Yan et al., 2008), mated to WT, unbarcoded *S. cerevisiae*. Fitness data from (Payen et al 2016). |
| Hybrid Barcoded Control Library | Collection of barcoded WT haploid *S. cerevisiae* strains from (Yan et al., 2008) mated to WT *S. uvarum*. |

### Fitness effects of hemizygosity in S. cerevisiae diploids

Both collections of control WT strains have narrow fitness distributions around neutrality, with 98% of the *S. cerevisiae* controls falling between fitness values of 0.047 and -0.040; and 98% of hybrid controls between 0.046 and -0.032 across all experiments (Supplemental Fig. S3). We used these empirical 1% cutoffs to determine significant increases and decreases in fitness of the deletion strains. Out of a total of 6,003 possible deletion strains, we identified 4,806 strains by barcode sequencing in the glucose-limited competition, 4,855 strains in phosphate limitation, and 4,901 strains in sulfate limitation. Compared to the WT control distribution, hemizygous gene deletions in *S. cerevisiae* caused a broader distribution of fitness effects. The null expectation for 1% cutoffs would be 48, 49, and 49 outliers in each direction for glucose, phosphate, and sulfate limitations, respectively. We observe significantly more deletion strains with fitness values beyond our cutoffs in several conditions: 308 haploinsufficient genes and 64 haploproficient genes in glucose limitation (p<2.2*10^-16^, p=0.19); 163 and 5 in phosphate limitation (p=4.3*10^-15^, p=4.4*10^-9^ fewer); and 58 and 113 in sulfate limitation (p=0.44, p=6*10^-7^; Comparison of Two Population Proportions performed in R, with Yates continuity correction). Thus we conclude that in in the *S. cerevisiae* hemizygous collection, there are more deletions that cause extreme fitness effects than we would expect by chance, consistent with our previous results (Payen et al. 2016).

### Fitness effects of LOH are more extreme in hybrids

We applied this analysis to the hybrid deletion strains. Out of 4,828 possible deletion strains (a lower number than above because only nonessential *S. cerevisiae* gene deletions can be used), we identified 3,195 deletion strains in sulfate limitation, 3,179 in phosphate limitation, and 2,955 in glucose limitation. Under the null expectation of 1%, we would expect 32 outliers in each direction in sulfate limited culture, 32 in phosphate limited, and 30 in glucose limited. However, in the hybrids we observed 308 haploinsufficient genes and 63 haploproficient genes in glucose limitation (p<2.2*10^-16^, p=0.0008); 919 and 453 in phosphate limitation (p<2.2*10^-16^, p<2.2*10^-16^); and 216 and 17 in sulfate limitation (p<2.2 * 10^-16^, p>0.05; Comparison of Two Population Proportions performed in R, with Yates continuity correction). The differences between nutrient limitations are illustrated in the different shapes of the distributions (Supplemental Figure S3). In phosphate limitation, the highest fitness strains had risen to an abundance >1.5% of the population by the final time point, over two orders of magnitude above their initial frequency.

Hybrid deletion mutants had a significantly broader range of fitness values than deletions of the same loci in the *S. cerevisiae* context (Supplemental Figure S3; Levene test p=1.9 * 10^-12^, p< 2.2*10^-16^, p=8.6*10^-6^, for glucose, phosphate, and sulfate limitations respectively) suggesting loss of one allele in hybrids leads to more extreme fitness outcomes, in both directions. We next compared genes that had significant fitness effects in hybrids to those in the *S. cerevisiae* diploids. Although there were some genes that were consistent between genetic backgrounds, correlation was low, and some gene deletions even had inverse effects (Fig. 1). Consistent with these findings, the genes identified in the hybrid and *S. cerevisiae* diploid datasets had different GO enrichments (Supplemental Tables S3 and S4).

**Figure 1.**
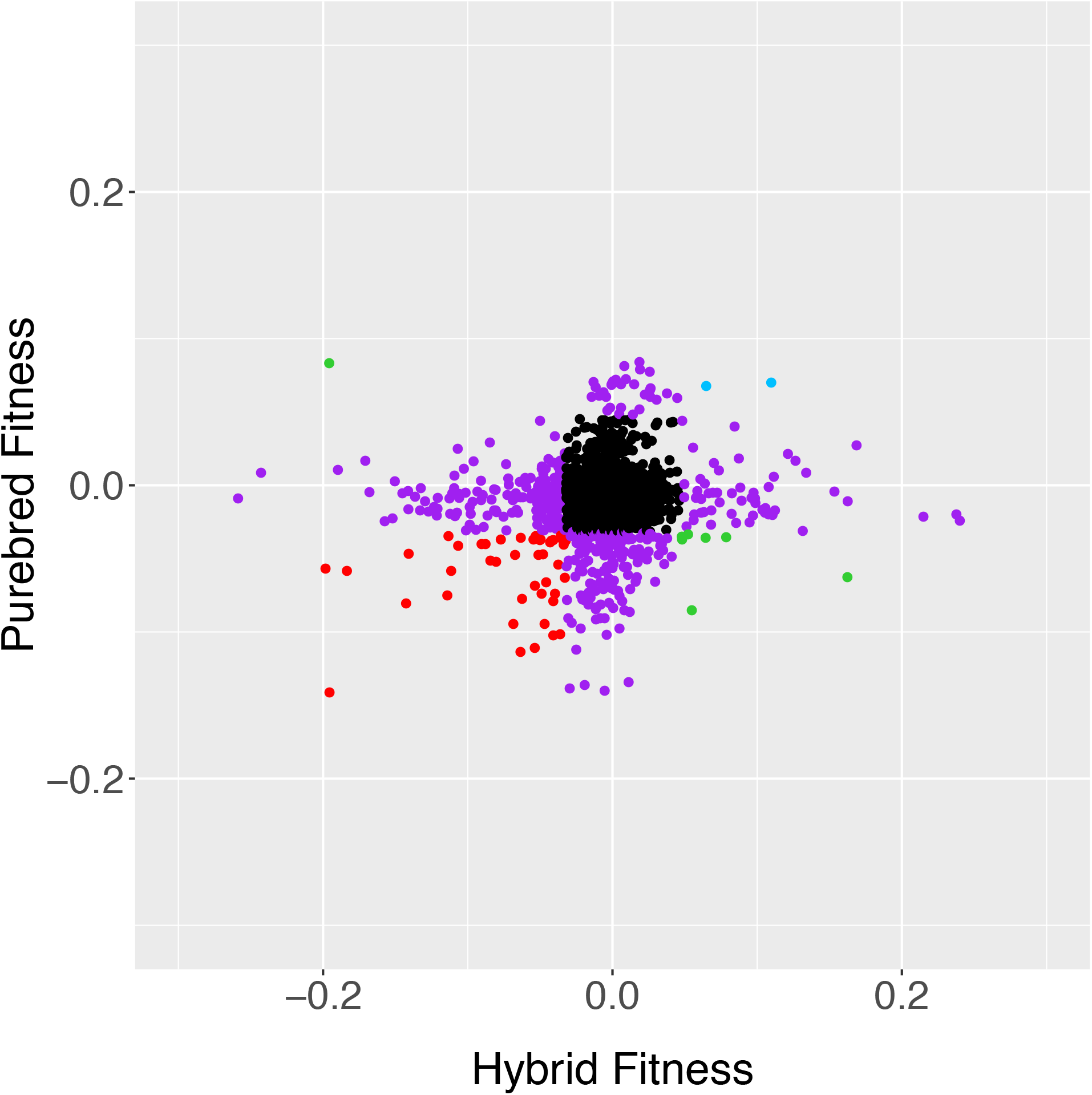
Scatter plot of fitness values for hybrid and diploid *S. cerevisiae* strains hemizygous for deletion mutations, measured in glucose limitation (comparisons in other nutrient limitations can be found in Supplemental Figure S4). Black strains fall inside the 1% cutoff in both axes, purple strains fall outside the 1% cutoff in just one axis, and the other colors fall outside of the cutoffs in both axes. Data from Supplemental Table S1. R^2^ = 0.00

Highlighting the effect of genetic background, many of the haploinsufficient alleles in the purebred genetic context are alleviated in the hybrid context, defined as an increase in fitness of at least 0.04, which is the 95% cutoff for the purebred collection. In glucose, phosphate, and sulfate limitations there are 93, 54 and 44 such alleviations of haploinsufficiency, respectively. This represents an alleviation rate of 30% and 34% in phosphate and glucose limited media, and 76% alleviation in sulfate limited media. GO enrichments for these alleviated gene deletions include cytosolic ribosomal subunit and ubiquinone metabolic process in glucose limitation; retrograde transport, endosome to golgi, and large ribosomal subunit for phosphate limitation; and the core mediator complex in sulfate limitation (all p-values<0.05 with a Bonferroni step down correction).

### Fitness effects of LOH are condition-specific

We next looked across environments to determine if hemizygosity caused larger fitness differences between conditions, or if effects were condition-specific. We calculated the variance in fitness values across conditions of 2,775 gene deletions present in all 6 mass competitions (hybrid and purebred deletion collections completed in the 3 nutrient limitations). Hybrid deletion strains had significantly larger variance between conditions relative to their purebred counterparts (T-test p<2.2*10^-16^). In the hybrid genetic context, 92 deletion strains showed antagonistic pleiotropy—low fitness in one condition and high fitness in another. These genes were enriched for GO terms gene expression and RNA metabolic process (p=2.5*10^-6^, and p=4.3*10^-6^), suggesting that differential expression may contribute to this pleiotropic phenotype.

One consequence of these patterns is that fitness in one nutrient limitation did not predict fitness in the others (Fig. 2). No loci showed consistent fitness differences across all three environments and in both genetic backgrounds. However, 89 deletions caused fitness deficits in two of the media, with no effect in the third (Supplemental Table S5), and 22 genes were haploproficient in two media and neutral in the third (Table 2). These genes are of particular interest because they may allow hybrid strains to adapt to multiple or heterogeneous environments. We previous found mutations in one of these genes, *MHR1*, in two phosphate-limited evolved populations (Smukowski Heil et al. 2017), showing the efficacy of this approach in finding potential driver mutations.

**Figure 2.**
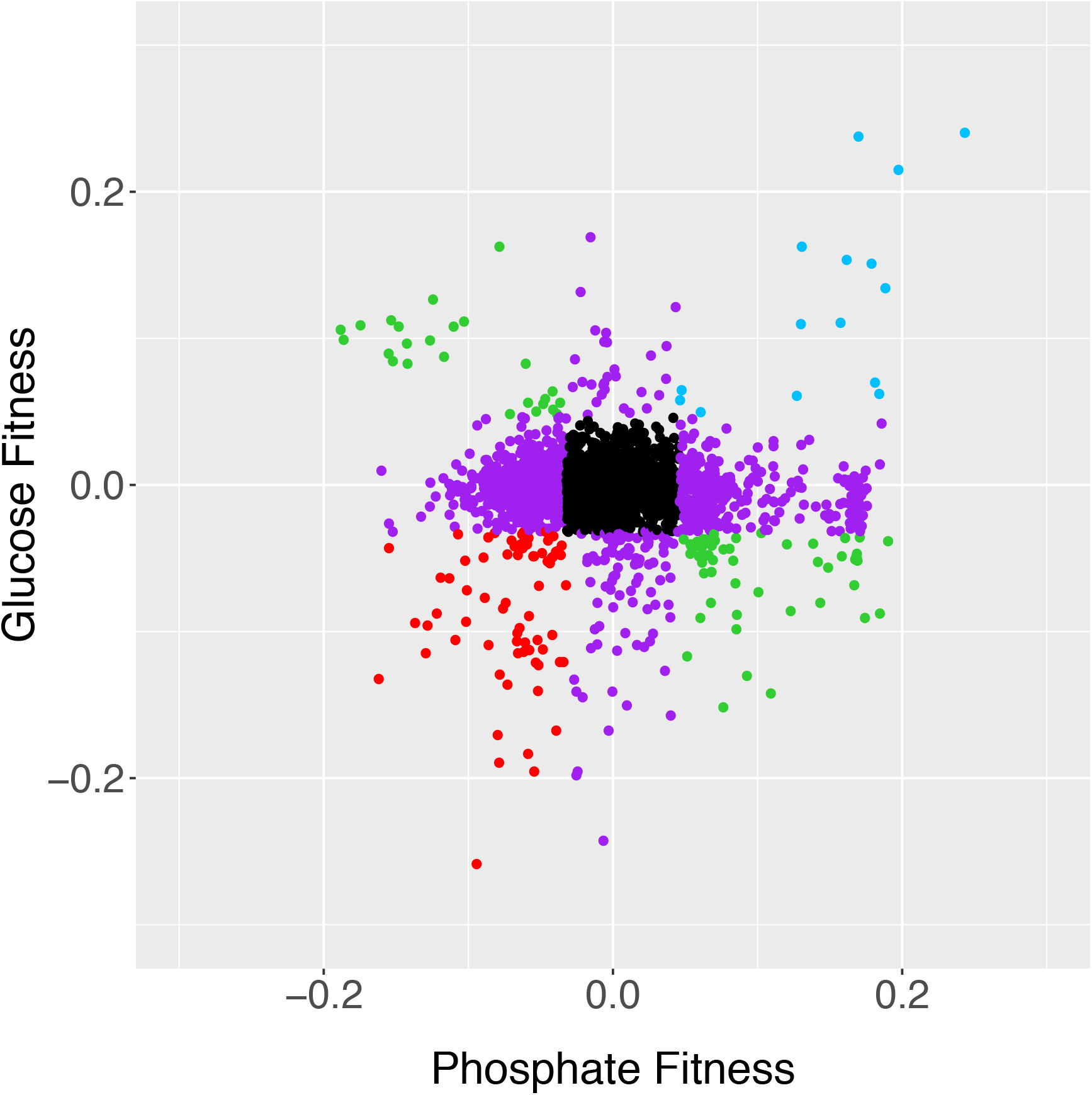
Fitness values of hybrids compared in glucose and phosphate limitation. Strains in black fall inside the 1% cutoff in both axes, purple strains fall outside the 1% cutoff in just one axis, and the other colors fall outside of the cutoffs in both axes. Comparisons between other media are shown in Supplemental Fig. S5. Data from Supplemental Table S1. R^2^=0.00

**Table 2.**
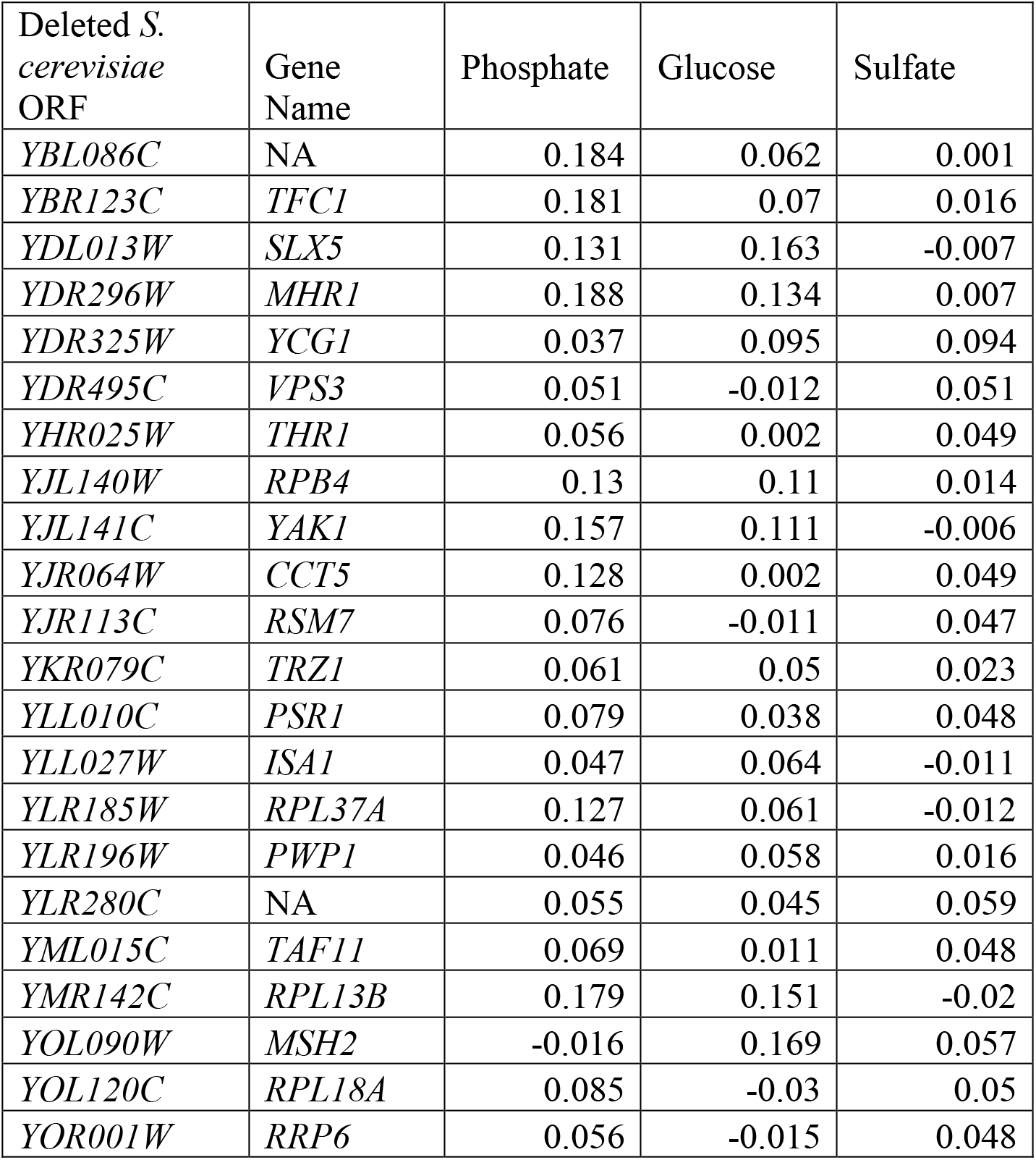
Condition-dependent haploproficiency in hybrid yeast. The last three columns are the fitness values of hemizygous deletion strains grown in the labeled limitation.

| Deleted *S. cerevisiae* ORF | Gene Name | Phosphate | Glucose | Sulfate |
| --- | --- | --- | --- | --- |
| *YBL086C* | NA | 0.184 | 0.062 | 0.001 |
| *YBR123C* | *TFC1* | 0.181 | 0.07 | 0.016 |
| *YDL013W* | *SLX5* | 0.131 | 0.163 | -0.007 |
| *YDR296W* | *MHR1* | 0.188 | 0.134 | 0.007 |
| *YDR325W* | *YCG1* | 0.037 | 0.095 | 0.094 |
| *YDR495C* | *VPS3* | 0.051 | -0.012 | 0.051 |
| *YHR025W* | *THR1* | 0.056 | 0.002 | 0.049 |
| *YJL140W* | *RPB4* | 0.13 | 0.11 | 0.014 |
| *YJL141C* | *YAK1* | 0.157 | 0.111 | -0.006 |
| *YJR064W* | *CCT5* | 0.128 | 0.002 | 0.049 |
| *YJR113C* | *RSM7* | 0.076 | -0.011 | 0.047 |
| *YKR079C* | *TRZ1* | 0.061 | 0.05 | 0.023 |
| *YLL010C* | *PSR1* | 0.079 | 0.038 | 0.048 |
| *YLL027W* | *ISA1* | 0.047 | 0.064 | -0.011 |
| *YLR185W* | *RPL37A* | 0.127 | 0.061 | -0.012 |
| *YLR196W* | *PWP1* | 0.046 | 0.058 | 0.016 |
| *YLR280C* | NA | 0.055 | 0.045 | 0.059 |
| *YML015C* | *TAF11* | 0.069 | 0.011 | 0.048 |
| *YMR142C* | *RPL13B* | 0.179 | 0.151 | -0.02 |
| *YOL090W* | *MSH2* | -0.016 | 0.169 | 0.057 |
| *YOL120C* | *RPL18A* | 0.085 | -0.03 | 0.05 |
| *YOR001W* | *RRP6* | 0.056 | -0.015 | 0.048 |

### Fitness effects of LOH are allele-specific

In our previous work, we found that loss of heterozygosity at the *PHO84* locus was beneficial when either allele was lost, i.e. heterozygosity itself had a cost (Smukowski Heil et al., 2017). To determine whether this phenomenon is widespread in our genome-scale dataset, we performed reciprocal hemizygosity analyses (Steinmetz et al., 2002). We deleted 11 *S. uvarum* genes that represented a broad range of fitness values (Supplemental Table S6), and mated these strains to WT *S. cerevisiae*, creating a set of reciprocal deletion strains vs. our original experiment. We then competed each strain against a WT hybrid labeled with GFP in the indicated nutrient limitation. With one exception (*TPK3*), the fitness values of these experiments were uncorrelated with those obtained with the corresponding *S. cerevisiae* allele deleted (Supplemental Table 6; Fig. 3).

**Figure 3.**
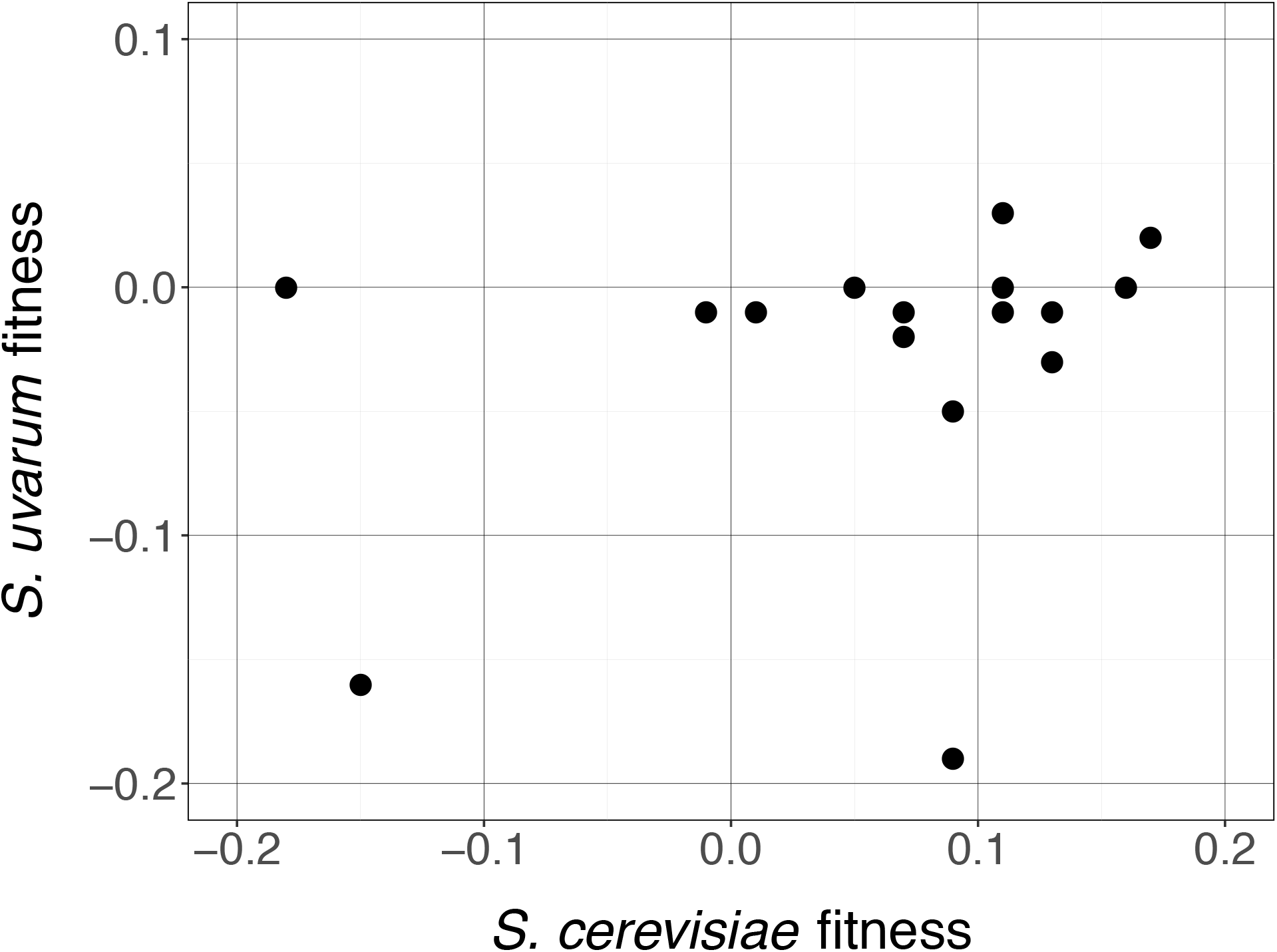
Reciprocal hemizygosity analysis correlation between *S. cerevisiae* deletion (X-axis) and *S. uvarum* deletion (Y-axis). R^2^=0.09, p=0.25. All nutrient environments included.

### Candidate genes driving LOH in experimental evolution

Together, these results show that beneficial mutations in hybrids cannot be predicted on the basis of screens performed in *S. cerevisiae* alone. LOH events may be tens of kilobases long and include hundreds of genes, making it impractical to use single gene approaches to understand such events. Instead, we used this genome-wide hybrid screen to identify candidate beneficial mutations implicated in LOH events from evolved strains (Smukowski Heil et al. 2017). Six out of 16 hybrid strains contained a total of nine LOH regions, and four of these regions eliminated the *S. cerevisiae* portion of the genome. These strains have fitness benefits compared to the fully heterozygous ancestor strain. One strain containing two LOH regions was evolved in sulfate limitation, where adaptation is largely dominated by amplification of the *SUL1* sulfate transporter gene (Brewer et al. 2015; Sanchez et al. 2017a). In this strain, we did not identify any single candidate gene deletions that had fitness benefits above our control strain cutoff of 0.046 in either region, consistent with this hypothesis. However, in phosphate limitation, we found candidate driver mutations for the regions deleted in both evolved strains (Table 3). Similar to our results for the whole dataset, none of these candidate gene deletions were beneficial in other nutrient limitations in the hybrid context, or in any nutrient limitation in the *S. cerevisiae* diploid. They also spanned a diverse variety of biological processes, including a gene of unknown function.

**Table 3.** Candidate driver genes in deleted *S. cerevisiae* genomic segments in evolved phosphate-limited hybrid strains.

| Candidate driver mutation<br>(systematic gene name of *S. cerevisiae* deleted gene) | Gene name | Competitive fitness in<br>phosphate limitation from<br>hemizygous hybrid screen |
| --- | --- | --- |
| *YIR023W* | *DAL81* | 0.132 |
| *YML041C* | *VPS71* | 0.080 |
| *YML049C* | *RSE1* | 0.171 |
| *YML061C* | *PIF1* | 0.048 |
| *YML066C* | *SMA2* | 0.072 |
| *YML069W* | *POB3* | 0.055 |
| *YML091C* | *RPM2* | 0.067 |
| *YML105C* | *SEC65* | 0.068 |
| *YML112W* | *CTK3* | 0.169 |
| *YML115C* | *VAN1* | 0.168 |
| *YML128C* | *MSC1* | 0.131 |
| *YML130C* | *ERO1* | 0.061 |
| *YML131W* | NA | 0.051 |

## DISCUSSION

LOH is prevalent in hybrid genomes across taxa. In our previous work, we observed these events arising quickly in interspecific yeast hybrids during only a few hundred generations of laboratory selection (Smukowski Heil et al. 2017; 2018b). In this study, we found specific gene deletions within these regions that might contribute to the fitness benefits enjoyed by these evolved strains, and we broadened our analysis to the whole genome to determine how hemizygous deletions behave more generally. We find that hybrids are more likely than purebred diploids to benefit from hemizygous deletions (but also to suffer fitness penalties). However, these benefits are complex – hemizygous deletions are largely condition-, allele-, and species-specific. Our results suggest that LOH may be an attractive means by which hybrids can adapt to strong, narrow selection pressures, but at the cost of reduced fitness in alternate environments. Industrial and fermentation environments, where yeast hybrids are successful, might provide exactly such a scenario (Hittinger 2013; Krogerus et al. 2017; Krogerus et al. 2018).

Though we have focused here on beneficial gene deletions, we note that other groups have used similar datasets to look at mutations that decrease fitness (Herbst et al. 2017; Weiss et al. 2018). We largely confirm the patterns observed in *S. paradoxus* x *S. cerevisiae* hybrids, though the different conditions utilized across studies make direct comparisons complicated.

We note several points for improvement and further study. First, while we did observe some significant gene ontology enrichments among gene deletions with shared fitness characteristics (for example differences in ribosomal structure/function and expression differences may contribute to differential fitness in various genetic and environmental contexts, respectively), the lack of strong biological process enrichments provided no simple rationale for the molecular explanations underlying these effects. Combining our data with other results collected from hybrids, such as protein-protein interactions (Chretien et al. 2018) may help provide such explanations. Expanding our study to additional genes would also be desirable. Due to the method we used to generate the hybrid deletion strains, we were limited to genes that are nonessential in *S. cerevisiae*. Though we deleted a small number of *S. uvarum* genes to compare with the orthologous *S. cerevisiae* allele deletions, we did not explore *S. uvarum* genes genome wide. We hope to remedy this in the future using our recently created *S. uvarum* insertional mutagenesis library (Sanchez et al. 2017b). *In vivo* transposition is another potential approach that has recently been applied to *S. paradoxus* x *S. cerevisiae* hybrids (Weiss et al. 2018). Both these approaches even have some advantages over the *S. cerevisiae* deletion collection, which has potential problems with suppressor mutations, mutation accumulation, and aneuploidy (Hughes et al. 2000; Teng et al. 2013; van Leeuwen et al. 2016).

Finally, we have not been able to determine how multiple genes within a segment combine to generate their full fitness consequence, a topic that has bedeviled the aneuploidy field more broadly (Solimini et al. 2012; Davoli et al. 2013; Sunshine et al. 2015; Dodgson et al. 2016; Iyer et al. 2018). Comparison of the fitness values of the evolved hybrids with the fitness of single gene deletions revealed cases where a simple additive model both under- or over-estimated the evolved strain fitness (analysis not shown; data in Table 2 and in Smukowski Heil, et al, 2017). However, the evolved strains are an imperfect basis for comparison since they contain other mutations in addition to the hemizygous region. Alternative selection methods for recovering high fitness strains with a minimal number of additional mutations presents one possible approach (Bellon et al. 2018). An even better approach might be to create hybrids with segmental monosomy and test their fitness directly. Such strains could be engineered using chromosome fragmentation vectors or recombinase-based approaches such as the Sc2.0 shuffle system (Morrow et al. 1997; Dymond et al. 2011; Shen et al. 2018).

## METHODS

### Strains and collections

The *S. cerevisiae* strain collections are described in Payen et al, 2016 (Payen et al. 2016).

All *S. uvarum* strains are derived from the reference strain CBS 7001, sometimes identified as *S. bayanus* var *uvarum*.

Hybrid collections were made by spreading 200 μl mid log *S. uvarum lys2 MAT*α on solid YPD omni plates then spotting the appropriate haploid *MAT***a** collection (deletion or barcoder) using a pinner with 96 arrayed pins. After overnight growth, colonies were transferred to selective solid minimal media to ensure only hybrid growth. These plates were then transferred to liquid selective media and pooled.

*S. uvarum* deletion strains were provided by Sarah Bissonnette and Jasper Rine (UC Berkeley) (Sanchez et al. 2017b).

### Fitness assays and barcode sequencing

All collections were grown as pools in duplicate in three different chemostat media – glucose limited, phosphate limited, and sulfate limited. All *S. cerevisiae* competition experiments were taken from Payen et al., 2016, where the protocol is described in detail. Briefly, pools were inoculated into 240 ml chemostats and grown for 24 hours, when peristaltic pumps were turned on at a dilution rate of ~0.17 volumes per hour (Payen et al. 2016). Samples were taken from chemostats twice a day. From these samples, the unique DNA barcodes from each collection were PCR amplified, with each time point having a unique Illumina adapter incorporated during PCR amplification. The barcodes were then sequenced on an Illumina Genome Analyzer IIx. The frequency of the barcodes was used to calculate the fitness of each strain by determining the natural log in the change of proportional barcode frequency over 25 generations. We required a minimum of 100 barcode counts per strain to be identified. For the reciprocal hemizygosity analysis, the 100 barcode limit was reduced to 44 for comparison to the mass competitions because manual curating of these ensured no false positives. Many sequences for the WT barcoded collection DNA barcodes were only determinable through examination of over represented sequences in sequencing data, and these were used for analysis.

Individual competition experiments were done in the respective media in 20 mL chemostats and competed against a single WT clone with a GFP label. Relative strain abundance was monitored using a BD Accuri C6 flow cytometer. Fitness was determined by regressing the slope of generations versus the ln(dark cells/GFP cells).

### GO Enrichments in the Dataset

GO enrichments were determined using the ClueGO application in Cytoscape (Bindea et al. 2009), and using the total strains identified in our experiments as the background population. Outliers were determined using a 1% cutoff in each direction based on the WT barcoded collection. All ontologies were corrected for multiple comparisons with a Bonferroni step down analysis.

### Statistics

Statistical measures unless otherwise stated were performed in R. The statistics used are stated in the results adjacent to p-values.

## DATA ACCESS

Sequencing data are available via BioProject Accession PRJNA283983.

## ACKNOWLEDGEMENTS

Thanks to Caitlin Connelly for her contributions to the initiation of this project. Thanks to Corey Nislow, Jasper Rine, and Sarah Bissonnette for sharing strains.

## SUPPLEMENTAL TABLE AND FIG. LEGENDS

Supplemental Table 1. Mass competitive fitness values of all deletion strains in our dataset across both species and all three nutrient limitations. NAs are samples for which there are no fitness values.

Supplemental Table 2. Comparison of fitness values determined with *en masse* competition experiments to fitness values determined in individual competition experiments.

Supplemental Table 3. GO results determined using Cytoscape in the purebred hemizygous deletion population.

Supplemental Table 4. GO results determined using Cytoscape in the hybrid hemizygous deletion population.

Supplemental Table 5. Haploinsufficient loci in at least two different nutrient limitations, as determined by distribution of wild-type fitness values, in the hybrid genetic background.

Supplemental Table 6. Reciprocal hemizygosity analysis. Fitness of hybrid strains with deleted *S. uvarum* genes and homologous deleted *S. cerevisiae* genes.

Supplemental Figure 1. Comparisons of fitness values derived from *en masse* hybrid hemizygote replicates in glucose, phosphate, and sulfate limitations.

Supplemental Figure 2. Comparison of fitness of hemizygous mutants between *en masse* competition experiments and individual competition experiments.

Supplemental Figure 3. Fitness distribution of WT hybrids and hemizygous hybrids (B, D, F), and WT purebreds and hemizygous purebreds (A,C,E). Green is the hemizygous deletion distribution and blue is the WT distribution of fitness in (A,B) glucose, (C,D) phosphate and (E,F) sulfate limited media.

Supplemental Figure 4. Comparison between hybrid and purebred fitness in A) phosphate and B) sulfate limitations. Black strains fall inside the 1% cutoff in both axes, purple strains fall outside the 1% cutoff in just one axis, and the other colors fall outside of the cutoffs in both axes. Both R^2^ = 0.00.

Supplemental Figure 5. Comparison of hybrid hemizygous fitness values between A) sulfate and phosphate limitations and B) glucose and sulfate limitations. Black strains fall inside the 1% cutoff in both axes, purple strains fall outside the 1% cutoff in just one axis, and the other colors fall outside of the cutoffs in both axes. Both R^2^ = 0.00.

## REFERENCES

Almeida P, Goncalves C, Teixeira S, Libkind D, Bontrager M, Masneuf-Pomarede I, Albertin W, Durrens P, Sherman DJ, Marullo P et al. 2014. A Gondwanan imprint on global diversity and domestication of wine and cider yeast *Saccharomyces uvarum*. Nat Commun 5: 4044.

Bellon JR, Ford CM, Borneman AR, Chambers PJ. 2018. A Novel Approach to Isolating Improved Industrial Interspecific Wine Yeasts Using Chromosomal Mutations as Potential Markers for Increased Fitness. Front Microbiol 9: 1442.

Bindea G, Mlecnik B, Hackl H, Charoentong P, Tosolini M, Kirilovsky A, Fridman WH, Pages F, Trajanoski Z, Galon J. 2009. ClueGO: a Cytoscape plug-in to decipher functionally grouped gene ontology and pathway annotation networks. Bioinformatics 25: 1091–1093.

Borneman AR, Zeppel R, Chambers PJ, Curtin CD. 2014. Insights into the *Dekkera bruxellensis* genomic landscape: comparative genomics reveals variations in ploidy and nutrient utilisation potential amongst wine isolates. PLoS Genet 10: e1004161.

Brenchley R, Spannagl M, Pfeifer M, Barker GL, D’Amore R, Allen AM, McKenzie N, Kramer M, Kerhornou A, Bolser D et al. 2012. Analysis of the bread wheat genome using whole-genome shotgun sequencing. Nature 491: 705–710.

Brewer BJ, Payen C, Di Rienzi SC, Higgins MM, Ong G, Dunham MJ, Raghuraman MK. 2015. Origin-Dependent Inverted-Repeat Amplification: Tests of a Model for Inverted DNA Amplification. PLoS Genet 11: e1005699.

Chen J, Upadhyaya NM, Ortiz D, Sperschneider J, Li F, Bouton C, Breen S, Dong C, Xu B, Zhang X et al. 2017. Loss of *AvrSr50* by somatic exchange in stem rust leads to virulence for Sr50 resistance in wheat. Science 358: 1607–1610.

Chester M, Leitch AR, Soltis PS, Soltis DE. 2010. Review of the Application of Modern Cytogenetic Methods (FISH/GISH) to the Study of Reticulation (Polyploidy/Hybridisation). Genes (Basel) 1: 166–192.

Chretien AE, Gagnon-Arsenault I, Dube AK, Barbeau X, Despres PC, Lamothe C, Dion-Cote AM, Lague P, Landry CR. 2018. Extended linkers improve the detection of protein-protein interactions (PPIs) by dihydrofolate reductase protein-fragment complementation assay (DHFR PCA) in living cells. Mol Cell Proteomics 17: 549.

Csoma H, Zakany N, Capece A, Romano P, Sipiczki M. 2010. Biological diversity of *Saccharomyces* yeasts of spontaneously fermenting wines in four wine regions: comparative genotypic and phenotypic analysis. Int J Food Microbiol 140: 239–248.

Dannemann M, Andres AM, Kelso J. 2016. Introgression of Neandertal- and Denisovan-like Haplotypes Contributes to Adaptive Variation in Human Toll-like Receptors. Am J Hum Genet 98: 22–33.

Davoli T, Xu AW, Mengwasser KE, Sack LM, Yoon JC, Park PJ, Elledge SJ. 2013. Cumulative haploinsufficiency and triplosensitivity drive aneuploidy patterns and shape the cancer genome. Cell 155: 948–962.

Delneri D, Hoyle DC, Gkargkas K, Cross EJ, Rash B, Zeef L, Leong HS, Davey HM, Hayes A, Kell DB et al. 2008. Identification and characterization of high-flux-control genes of yeast through competition analyses in continuous cultures. Nat Genet 40: 113–117.

Deutschbauer AM, Davis RW. 2005. Quantitative trait loci mapped to single-nucleotide resolution in yeast. Nat Genet 37: 1333–1340.

Dodgson SE, Kim S, Costanzo M, Baryshnikova A, Morse DL, Kaiser CA, Boone C, Amon A. 2016. Chromosome-Specific and Global Effects of Aneuploidy in *Saccharomyces cerevisiae*. Genetics 202: 1395–1409.

Dunn B, Sherlock G. 2008. Reconstruction of the genome origins and evolution of the hybrid lager yeast *Saccharomyces pastorianus*. Genome Res 18: 1610–1623.

Dymond JS, Richardson SM, Coombes CE, Babatz T, Muller H, Annaluru N, Blake WJ, Schwerzmann JW, Dai J, Lindstrom DL et al. 2011. Synthetic chromosome arms function in yeast and generate phenotypic diversity by design. Nature 477: 471–476.

Emery M, Willis MMS, Hao Y, Barry K, Oakgrove K, Peng Y, Schmutz J, Lyons E, Pires JC, Edger PP et al. 2018. Preferential retention of genes from one parental genome after polyploidy illustrates the nature and scope of the genomic conflicts induced by hybridization. PLoS Genet 14: e1007267.

Giaever G, Chu AM, Ni L, Connelly C, Riles L, Veronneau S, Dow S, Lucau-Danila A, Anderson K, Andre B et al. 2002. Functional profiling of the *Saccharomyces cerevisiae* genome. Nature 418: 387–391.

Gittelman RM, Schraiber JG, Vernot B, Mikacenic C, Wurfel MM, Akey JM. 2016. Archaic Hominin Admixture Facilitated Adaptation to Out-of-Africa Environments. Curr Biol 26: 3375–3382.

Gonzalez SS, Barrio E, Gafner J, Querol A. 2006. Natural hybrids from *Saccharomyces cerevisiae*, *Saccharomyces bayanus* and *Saccharomyces kudriavzevii* in wine fermentations. FEMS Yeast Res 6: 1221–1234.

Gresham D, Desai MM, Tucker CM, Jenq HT, Pai DA, Ward A, DeSevo CG, Botstein D, Dunham MJ. 2008. The repertoire and dynamics of evolutionary adaptations to controlled nutrient-limited environments in yeast. PLoS Genet 4: e1000303.

Herbst RH, Bar-Zvi D, Reikhav S, Soifer I, Breker M, Jona G, Shimoni E, Schuldiner M, Levy AA, Barkai N. 2017. Heterosis as a consequence of regulatory incompatibility. BMC Biol 15: 38.

Hittinger CT. 2013. *Saccharomyces* diversity and evolution: a budding model genus. Trends Genet 29: 309–317.

Huerta-Sanchez E, Jin X, Asan, Bianba Z, Peter BM, Vinckenbosch N, Liang Y, Yi X, He M, Somel M et al. 2014. Altitude adaptation in Tibetans caused by introgression of Denisovan-like DNA. Nature 512: 194–197.

Hughes TR, Roberts CJ, Dai H, Jones AR, Meyer MR, Slade D, Burchard J, Dow S, Ward TR, Kidd MJ et al. 2000. Widespread aneuploidy revealed by DNA microarray expression profiling. Nat Genet 25: 333–337.

Iyer J, Singh MD, Jensen M, Patel P, Pizzo L, Huber E, Koerselman H, Weiner AT, Lepanto P, Vadodaria K et al. 2018. Pervasive genetic interactions modulate neurodevelopmental defects of the autism-associated 16.11.2 deletion in *Drosophila melanogaster*. Nat Commun. 9:2548.

Krogerus K, Holmstrom S, Gibson B. 2018. Enhanced Wort Fermentation with De Novo Lager Hybrids Adapted to High-Ethanol Environments. Appl Environ Microbiol 84.

Krogerus K, Magalhaes F, Vidgren V, Gibson B. 2017. Novel brewing yeast hybrids: creation and application. Appl Microbiol Biotechnol 101: 65–78.

Kunicka-Styczynska A, Rajkowska K. 2011. Physiological and genetic stability of hybrids of industrial wine yeasts *Saccharomyces sensu stricto* complex. J Appl Microbiol 110: 1538–1549.

Libkind D, Hittinger CT, Valerio E, Goncalves C, Dover J, Johnston M, Goncalves P, Sampaio JP. 2011. Microbe domestication and the identification of the wild genetic stock of lager-brewing yeast. Proc Natl Acad Sci U S A 108: 14539–14544.

Louis VL, Despons L, Friedrich A, Martin T, Durrens P, Casaregola S, Neuveglise C, Fairhead C, Marck C, Cruz JA et al. 2012. *Pichia sorbitophila*, an Interspecies Yeast Hybrid, Reveals Early Steps of Genome Resolution After Polyploidization. G3 (Bethesda) 2: 299–311.

Maclean CJ, Greig D. 2008. Prezygotic reproductive isolation between *Saccharomyces cerevisiae* and *Saccharomyces paradoxus*. BMC Evol Biol 8: 1.

Marcet-Houben M, Gabaldon T. 2015. Beyond the Whole-Genome Duplication: Phylogenetic Evidence for an Ancient Interspecies Hybridization in the Baker’s Yeast Lineage. PLoS Biol 13: e1002220.

Martini AV, Martini A. 1987. Three newly delimited species of *Saccharomyces sensu stricto*. Antonie Van Leeuwenhoek 53: 77–84.

Mertens S, Steensels J, Saels V, De Rouck G, Aerts G, Verstrepen KJ. 2015. A large set of newly created interspecific *Saccharomyces* hybrids increases aromatic diversity in lager beers. Appl Environ Microbiol 81: 8202–8214.

Mixao V, Gabaldon T. 2018. Hybridization and emergence of virulence in opportunistic human yeast pathogens. Yeast 35: 5–20.

Morales L, Dujon B. 2012. Evolutionary role of interspecies hybridization and genetic exchanges in yeasts. Microbiol Mol Biol Rev 76: 721–739.

Morrow DM, Connelly C, Hieter P. 1997. "Break copy" duplication: a model for chromosome fragment formation in *Saccharomyces cerevisiae*. Genetics 147: 371–382.

Muller LA, McCusker JH. 2009. A multispecies-based taxonomic microarray reveals interspecies hybridization and introgression in *Saccharomyces cerevisiae*. FEMS Yeast Res 9: 143–152.

Murphy HA, Zeyl CW. 2012. Prezygotic isolation between *Saccharomyces cerevisiae* and *Saccharomyces paradoxus* through differences in mating speed and germination timing. Evolution 66: 1196–1209.

Naumova ES, Naumov GI, Masneuf-Pomarede I, Aigle M, Dubourdieu D. 2005. Molecular genetic study of introgression between *Saccharomyces bayanus* and *S. cerevisiae*. Yeast 22: 1099–1115.

Nguyen HV, Legras JL, Neuveglise C, Gaillardin C. 2011. Deciphering the hybridisation history leading to the Lager lineage based on the mosaic genomes of *Saccharomyces bayanus* strains NBRC1948 and CBS380. PLoS One 6: e25821.

Ohnuki S, Ohya Y. 2018. High-dimensional single-cell phenotyping reveals extensive haploinsufficiency. PLoS Biol 16: e2005130.

Otto SP, Whitton J. 2000. Polyploid incidence and evolution. Annu Rev Genet 34: 401–437.

Payen C, Sunshine AB, Ong GT, Pogachar JL, Zhao W, Dunham MJ. 2016. High-Throughput Identification of Adaptive Mutations in Experimentally Evolved Yeast Populations. PLoS Genet 12: e1006339.

Perez-Traves L, Lopes CA, Querol A, Barrio E. 2014. On the complexity of the *Saccharomyces bayanus* taxon: hybridization and potential hybrid speciation. PLoS One 9: e93729.

Peris D, Lopes CA, Belloch C, Querol A, Barrio E. 2012. Comparative genomics among *Saccharomyces cerevisiae x Saccharomyces kudriavzevii* natural hybrid strains isolated from wine and beer reveals different origins. BMC Genomics 13: 407.

Peris D, Moriarty RV, Alexander WG, Baker E, Sylvester K, Sardi M, Langdon QK, Libkind D, Wang QM, Bai FY et al. 2017. Hybridization and adaptive evolution of diverse *Saccharomyces* species for cellulosic biofuel production. Biotechnol Biofuels 10: 78.

Piotrowski JS, Nagarajan S, Kroll E, Stanbery A, Chiotti KE, Kruckeberg AL, Dunn B, Sherlock G, Rosenzweig F. 2012. Different selective pressures lead to different genomic outcomes as newly-formed hybrid yeasts evolve. BMC Evol Biol 12: 46.

Pir P, Gutteridge A, Wu J, Rash B, Kell DB, Zhang N, Oliver SG. 2012. The genetic control of growth rate: a systems biology study in yeast. BMC Syst Biol 6: 4.

Pryszcz LP, Nemeth T, Saus E, Ksiezopolska E, Hegedusova E, Nosek J, Wolfe KH, Gacser A, Gabaldon T. 2015. The Genomic Aftermath of Hybridization in the Opportunistic Pathogen *Candida metapsilosis*. PLoS Genet 11: e1005626.

Racimo F, Gokhman D, Fumagalli M, Ko A, Hansen T, Moltke I, Albrechtsen A, Carmel L, Huerta-Sanchez E, Nielsen R. 2017. Archaic Adaptive Introgression in TBX15/WARS2. Mol Biol Evol 34: 509–524.

Sanchez MR, Miller AW, Liachko I, Sunshine AB, Lynch B, Huang M, Alcantara E, DeSevo CG, Pai DA, Tucker CM et al. 2017a. Differential paralog divergence modulates genome evolution across yeast species. PLoS Genet 13: e1006585.

Sanchez MR, Payen C, Cheong F, Hovde B, Bissonnette S, Arkin A, Skerker JM, Brem R, Caudy AA, Dunham MJ. 2017b. Transposon insertional mutagenesis in *Saccharomyces uvarum* reveals trans-acting effects influencing species dependent essential genes. bioRxiv doi:10.1101/218305.

Scannell DR, Frank AC, Conant GC, Byrne KP, Woolfit M, Wolfe KH. 2007. Independent sorting-out of thousands of duplicated gene pairs in two yeast species descended from a whole-genome duplication. Proc Natl Acad Sci U S A 104: 8397–8402.

Schroder MS, Martinez de San Vicente K, Prandini TH, Hammel S, Higgins DG, Bagagli E, Wolfe KH, Butler G. 2016. Multiple Origins of the Pathogenic Yeast *Candida orthopsilosis* by Separate Hybridizations between Two Parental Species. PLoS Genet 12: e1006404.

Shen MJ, Wu Y, Yang K, Li Y, Xu H, Zhang H, Li BZ, Li X, Xiao WH, Zhou X et al. 2018. Heterozygous diploid and interspecies SCRaMbLEing. Nat Commun 9: 1934.

Sipiczki M. 2008. Interspecies hybridization and recombination in *Saccharomyces* wine yeasts. FEMS Yeast Res 8: 996–1007.

Smukowski Heil C, Burton JN, Liachko I, Friedrich A, Hanson NA, Morris CL, Schacherer J, Shendure J, Thomas JH, Dunham MJ. 2018a. Identification of a novel interspecific hybrid yeast from a metagenomic spontaneously inoculated beer sample using Hi-C. Yeast 35: 71–84.

Smukowski Heil CS, DeSevo CG, Pai DA, Tucker CM, Hoang ML, Dunham MJ. 2017. Loss of Heterozygosity Drives Adaptation in Hybrid Yeast. Mol Biol Evol 34: 1596–1612.

Smukowski Heil C, Large CRL, Patterson K, Dunham MJ. 2018b. Temperature preference biases parental genome retention during hybrid evolution. biorXiv. doi: 10.1101/429803

Solimini NL, Xu Q, Mermel CH, Liang AC, Schlabach MR, Luo J, Burrows AE, Anselmo AN, Bredemeyer AL, Li MZ et al. 2012. Recurrent hemizygous deletions in cancers may optimize proliferative potential. Science 337: 104–109.

Stelkens RB, Brockhurst MA, Hurst GD, Miller EL, Greig D. 2014. The effect of hybrid transgression on environmental tolerance in experimental yeast crosses. J Evol Biol 27: 2507–2519.

Sun S, Xu J. 2009. Chromosomal rearrangements between serotype A and D strains in *Cryptococcus neoformans*. PLoS One 4: e5524.

Sunshine AB, Payen C, Ong GT, Liachko I, Tan KM, Dunham MJ. 2015. The fitness consequences of aneuploidy are driven by condition-dependent gene effects. PLoS Biol 13: e1002155.

Teng X, Dayhoff-Brannigan M, Cheng WC, Gilbert CE, Sing CN, Diny NL, Wheelan SJ, Dunham MJ, Boeke JD, Pineda FJ et al. 2013. Genome-wide consequences of deleting any single gene. Mol Cell 52: 485–494.

van Leeuwen J, Pons C, Mellor JC, Yamaguchi TN, Friesen H, Koschwanez J, Usaj MM, Pechlaner M, Takar M, Usaj M et al. 2016. Exploring genetic suppression interactions on a global scale. Science 354.

Weiss CV, Roop JI, Hackley RK, Chuong JN, Grigoriev IV, Arkin AP, Skerker JM, Brem RB. 2018. Genetic dissection of interspecific differences in yeast thermotolerance. Nat Genet doi:10.1038/s41588-018-0243-4.

